# Phenotypic Heterogeneity in the DNA Replication Stress Response Revealed by Quantitative Protein Dynamics Measurements

**DOI:** 10.1101/2022.06.08.495346

**Authors:** Brandon Ho, Raphael Loll-Krippleber, Nikko P. Torres, Andreas Cuny, Fabian Rudolf, Grant W. Brown

## Abstract

Cells respond to environmental stressors by activating programs that result in protein abundance and localization changes. The DNA damage and DNA replication stress responses have been heavily studied and provide exemplars of the roles of protein localization and abundance regulation in proper cellular stress response. While vast amounts of data have been collected to describe the dynamics of yeast proteins in response to numerous external stresses, few have assessed and compared both protein localization kinetics and phenotypic heterogeneity in the same context, particularly during DNA replication stress. We developed a robust yet simple quantification scheme to identify and measure protein localization change events (re-localization) and applied it to the 314 yeast proteins whose subcellular distribution changes following DNA replication stress. We captured different kinetics of protein re-localization, identified proteins with localization changes that were not detected in previous analyses, and defined the extent of heterogeneity in stress-induced protein re-localization. Our imaging platforms and analysis pipeline enables efficient measurements of protein localization phenotypes for single cells over time and will guide future work in elucidating the biological parameters that govern cellular heterogeneity.

## Introduction

Environmental changes and stresses trigger multiple intracellular events that affect protein function, including changes in transcriptional programs, regulation of protein abundance, and post-translational modifications, (Causton et al., 2001; Gasch, 2007; Gasch and Werner-Washburne, 2002; Gasch et al., 2000). In particular, regulated intracellular localization of proteins is a key factor that influences protein activity. Intuitively, a protein’s activity directly depends on its proximity to necessary cofactors and substrates. The ability of cells to control the localization of a protein provides an opportunity for regulation of protein function, and ultimately cellular function, that can both complement and supercede regulation of expression and post-translational modification. Systematic analysis of protein localization has revealed several key properties. Few proteins change in both abundance and localization simultaneously, indicating that expression and location are controlled differently (Chong et al., 2015; Mazumder et al., 2013; Tkach et al., 2012; Torres et al., 2016). The repatterning of proteins within cells is condition specific (Torres et al., 2016). Only a handful of proteins respond in at least five of the stress conditions examined to date (Torres and Brown, 2015), suggesting that protein localization changes reflect specific responses to particular environmental or genetic cues, rather than being a general stress response or an experimental artifact. Although protein localization has been systematically studied under conditions including DNA damage (Chong et al., 2015; Dénervaud et al., 2013; Tkach et al., 2012), oxidative stress and nutrient limitation (Breker et al., 2013), and in different genetic backgrounds (Chong et al., 2015), there is currently little insight into localization dynamics or regulation, owing in part to limitations of image analysis.

Systematic analyses of protein localization changes in eukaryotes typically take advantage of the proteome-scale collection of Saccharomyces cerevisiae strains expressing individual protein fusions to GFP (Huh et al., 2003). Most work-flows involve exposing the 5000 strains in the collection to a specific environmental or genetic perturbation, followed by imaging at one or several times after exposure. Early studies relied on manual inspection of images, whereas more recent analyses employed machine learning methods to predict protein localization (Kraus et al., 2017), with dynamics inferred from static images with the greatest time-lapse movie resolution, to our knowledge, of 20 minutes (Dénervaud et al., 2013). Automated analyses of different types have been applied to yeast protein localization screens to classify the location of each protein with respect to subcellular compartments, and to quantify the extent of protein localization within the given compartment (Chen et al., 2007a; Chong et al., 2015; Dénervaud et al., 2013; Kraus et al., 2017; Lu et al., 2018). Continued refinement of analysis methods has yielded increasing accuracy, but with some limitations. Analysis methods that focus on specifying protein origins and destinations are computationally intensive, require extensive hand-annotated training sets, and can fail to classify complicated re-localization patterns. Of all the environmental stress contexts, protein localization changes have been most thoroughly studied in the context of DNA damage and DNA replication stress (Chong et al., 2015; Dénervaud et al., 2013; Lisby et al., 2004; Mazumder et al., 2013; Tkach et al., 2012). The cellular response to DNA damage and DNA replication stress includes relocalizations, such as the concentration of proteins in punctate nuclear foci and transport from the cytoplasm to the nucleus, which have intimate mechanistic connections to stress resistance and to DNA repair. Interestingly, DNA damage- and DNA replication stress-induced relocalizations are not restricted to nuclear proteins, but span across all compartments of the cell (Chong et al., 2015; Dénervaud et al., 2013; Loll-Krippleber and Brown, 2017; Mazumder et al., 2013; Tkach et al., 2012). Protein re-localizations associated with DNA damage and DNA replication stress are also observed in metazoans, indicating broad use of protein localization regulation to control protein function, and suggesting that some regulatory modes could be shared by evolutionarily distant organisms. Here we developed an automated computational analysis to quantify changes in protein localization that is applicable to a wide array of fluorescence microscopic images. We applied our method to cells exposed to methyl methanesulfonate (MMS) and hydroxyurea (HU), imaging at seven time points over four hours, for an array of 314 proteins implicated in the replication stress response. We improved on existing data to expand the full repertoire of proteins that change localization during replication stress, and found that phenotypic penetrance for localization changes varies across the complete range of population fractions. Quantification of our images revealed diverse temporal responses of protein localization in replication stress, which we used to identify a new role of Mrx16 as a component of the intranuclear quality control (INQ) compartment. Finally, we imaged cells at high temporal resolution, demonstrating that replication stress evokes extensive single-cell heterogeneity in protein localization and that the factors governing this variation can be protein specific. Overall, our screening effort reveals new proteins involved in genome stability and quantifies the phenotypic heterogeneity that exists in the DNA replication stress response.

## Results

### Quantifying protein subcellular localization changes using intensity distribution

Based on the observation that the fluorescence intensity for any given green fluorescent protein (GFP)-tagged protein significantly increases when it is concentrated to a confined subcellular space, we reasoned that changes in the GFP pixel intensity distribution could be used as a proxy for protein localization. We also noted that localization changes can be detected independently of the exact nature of the localization itself, allowing solutions that are less computationally intense than quantifying the type of localization and the localization change simultaneously. In addition, if a method is blind to the type of localization then it could detect localizations that are not present in the image training set, and in fact would not require a training set. Finally, a quantification method that is independent of the type of localization is particularly suited for the yeast platform due to the extensive annotation of localizations already available in the literature (Chong et al., 2015; Huh et al., 2003; Mazumder et al., 2013; Tkach et al., 2012).

We developed a pipeline to measure changes in the GFP pixel intensity distribution following drug treatment (Figure 1a). The quantification scheme we describe captures general protein re-localization, irrespective of the direction of movement or type of subcellular change. Our strategy first segments and isolates single cells in our images then extracts the GFP signal intensity for each pixel within each cell. The GFP pixel intensities for these single cells is then normalized to the median GFP intensity of each respective cell to account for any total intensity changes that may occur during drug treatment, thereby minimizing the effect of changes in protein abundance on quantification of localization. Finally, the brightest pixel intensities are calculated for each cell as the 95th percentile of the normalized intensity distribution, hereafter referred to as the localization (LOC) value. As a protein concentrates into a particular subcellular space, the LOC value for that cell increases. Changes in LOC values over time can then be used to identify protein localization change events. At any given time within the experiment, if a cellular LOC value was two median absolute deviations (MAD) greater than or less than the median LOC values of all untreated cells (x +/- 2 MAD) the cell was labeled as having increased or decreased protein localization, respectively. Finally, the percentage of cells with a localization change was calculated for each timepoint of drug-treatment (Figure 1a, iv). A protein was considered to display a localization change if the maximum percent of cells exhibiting protein relocalization changed by at least 2-fold compared to the untreated population of cells.

**Figure 1:**
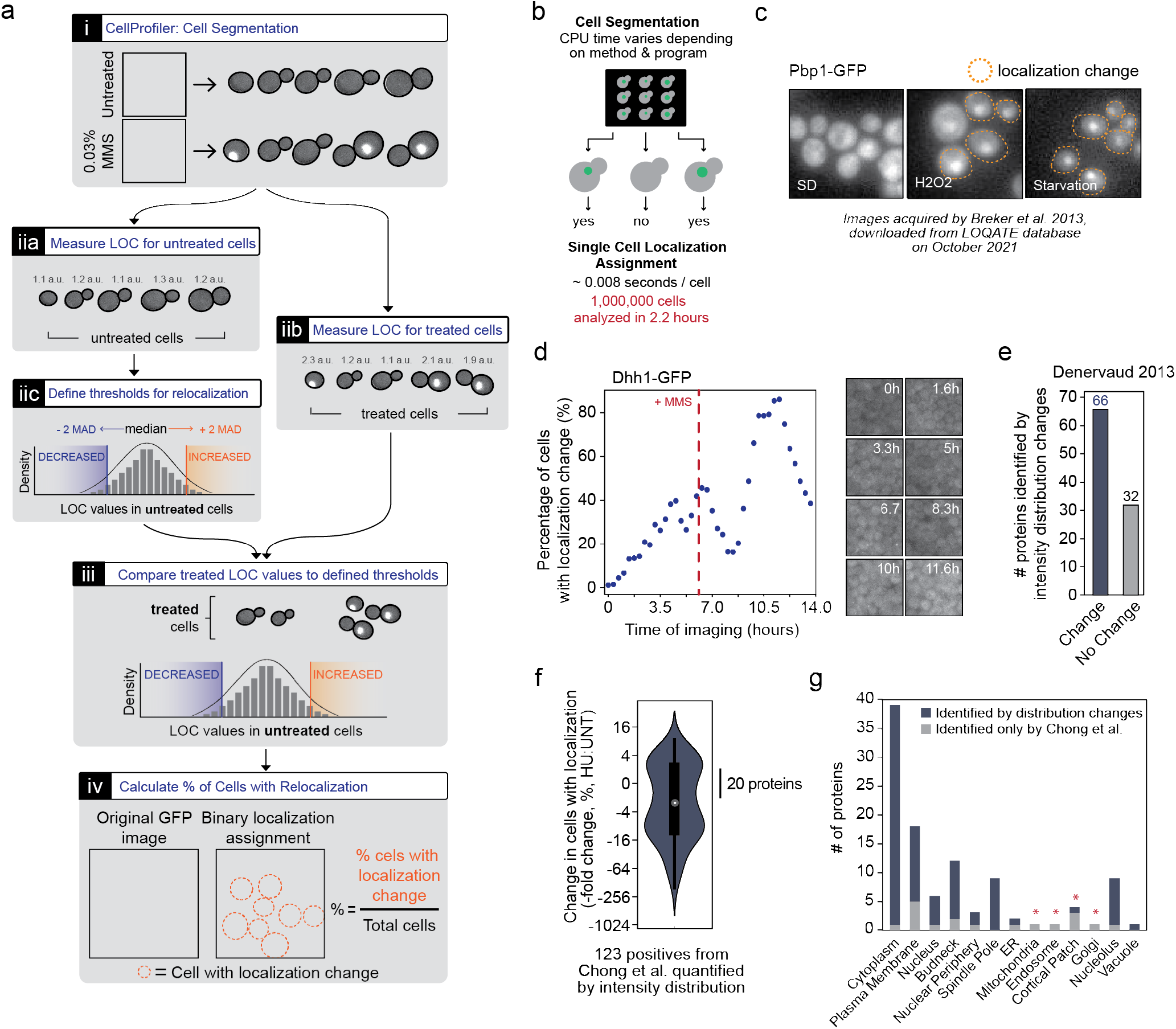
Protein localization changes are captured by pixel intensity distribution changes. (a) Quantification pipeline to measure protein re-localization. (i) Cells are segmented, (ii) the pixel intensity distribution for each individual cell is measured, referred to as the LOC score, and compared to the distribution for all untreated cells with a given protein-GFP. (iii) If the distribution changes and passes thresholds defined by the untreated cells, then the cell is labeled as harbouring a protein-GFP protein that has ‘re-localized’. (iv) The percent of cells with a protein re-localization is then calculated. (b) Schematic representing the analysis pipeline workflow, highlighting the two most computationally heavy tasks; cell segmentation and single-cell quantification (c) Representative images of Pbp1-GFP re-localization during peroxide and starvation stress, as acquired in (Breker et al., 2013, 2014). Cells identified as re-localized by our pipeline are outlined in orange. (d) My analysis pipeline was applied to images acquired in (Dénervaud et al., 2013) for 98 selected proteins. The quantification of Dhh1-GFP protein re-localization by our pipeline, and representative images of Dhh1-GFP cytoplasmic foci formation following 0.1% MMS treatment, are shown. (e) The number of proteins that we identify to change in protein localization (> 2-fold change, blue bar) from the set of proteins identified to change localization by (Dénervaud et al., 2013). (f) Images of all 123 proteins identified as changing in 160 mM HU from (Chong et al., 2015) were downloaded from CYCLoPS (Koh et al., 2015) and quantified by our pipeline. The log2 fold-change for each of these proteins is plotted as a violin plot. The grey dot represents the median and large black rectangle represents the interquartile range (25th-75th percentile). The orange and blue shaded background represent fold-change greater or less than two fold, respectively. (g) The subcellular compartments for all 123 proteins are plotted, and the proportion of these proteins our pipeline can detect as changing in localization is shown in blue. Red asterisks indicate subcellular compartments where Chong et al. perform better with regards to detection of localization.

Improvements in microscopic imaging throughput has greatly increased the need for effective strategies to measure cellular parameters on a large scale. Previous machine learning methods in outlier detection and pattern recognition have provided quantitative measurements of yeast protein localization with high accuracy (> 70% precision and recall) (Grys et al., 2017). Typically, these techniques can be difficult to apply to diverse datasets. As a result, users may be required to re-train published analysis methodologies to suit their images, which can be a challenging and time-consuming process. An unsupervised methodology has been designed to detect protein-GFP localization changes without any training (Lu and Moses, 2016; Lu et al., 2018). However, this method still requires the extraction of up to 120 single cell features and assembling feature vectors for each protein for comparative analyses with other literature data sets (Lu and Moses, 2016). Although our localization analysis reduces highly dimensional single-cell imaging data into a single measured parameter, our method is capable of quantifying protein localization changes rapidly without the need for imaging data other than those acquired by the user, distilling measurements from a million segmented single cells in just over two hours, with the limiting step in our quantification pipeline being single cell segmentation (Figure 1b).

A critical component of our analysis pipeline that makes it suitable for large-scale imaging data is its ability to identify localization changes in micrographs from a wide range of microscopic instruments and conditions. We tested the versatility of our pipeline by applying it to images of yeast cells acquired by diverse platforms. First, we focused on 16 randomly selected proteins annotated as changing localization in DTT, oxidative, and/or starvation stress conditions (Breker et al., 2013, 2014). We downloaded low-resolution vignette images from the LoQAtE database and processed them with our intensity distribution method. Despite the images often containing fewer than 12 cells, we found that 13 proteins out of 16 showed an increase in the fraction of cells displaying re-localization (Figure 1c). In a second dataset, cells were exposed to 0.1% MMS in a microfluidic device and wide-field fluorescence images (Nikon Eclipse Ti-E, Nikon Instruments) were acquired by Denervaud et al. (Dénervaud et al., 2013). They identified 118 proteins that changed localization in MMS conditions. We were able to download time-lapse images for 98 of these 118 proteins from CELLBASE (Dénervaud et al., 2013), to which we applied our analysis method (Figure 1d). We identified 66 out of 98 proteins as changing in localization, indicating good agreement across imaging platforms (Figure 1e).

Finally, we compared our pipeline to machine learning classifiers used to score images acquired on the same high-throughput confocal platform as our images (OPERA, PerkinElmer) in the HU stress condition screened by Chong et al. (Chong et al., 2015). At the level of binary re-localization calls for cell populations, agreement between the methods was excellent (84% of re-localizations noted by Chong et al were detected by our method). There were 20 instances where machine learning detected localization changes whereas intensity distribution did not (Figure 1e, 1f). Localization changes involving the mitochondria, Golgi, endosome, and cortical patches (Figure 1g) displayed the greatest discrepancy between our quantification and that of Chong et al, highlighting that machine learning methods are capable of greater sensitivity for proteins that exhibit complex localization patterns. At the more granular level of detecting single cells displaying a localization change there was some differences between the methods. Even among proteins with a localization change identified by the two methods, there are instances where Chong et al detects a greater change in localization. This is largely due to these proteins exhibiting high fluorescence signal and/or high localization in the untreated condition (Figure EV1), decreasing the dynamic range for intensity distributions to be detected by our method, resulting in an underestimation of localization changes. Nevertheless, we demonstrate that simple changes in GFP pixel intensity distributions can be a powerful tool for monitoring changes in proteome spatial dynamics across all subcellular compartments, with savings in compute time, and ease of comparison across image acquisition platforms.

### Intensity distribution analysis detects previously uncharacterized localization responses

We, and others, have identified several hundred proteins that change in intracellular location after chemically induced DNA replication stress imposed by the ribonucleotide reductase inhibitor HU or by the alkylating agent MMS (Chong et al., 2015; Dénervaud et al., 2013; Mazumder et al., 2013; Tkach et al., 2012). Combining high-throughput results with those from smallscale analyses, we generated a list of 314 proteins that change location during replication stress, to explore protein re-localization properties over longer durations of stress (Table EV1). Since DNA damage-induced sumoylation contributes to faithful replication and repair under DNA damage and replication stress conditions, this list of proteins also included identified sumoylated proteins targets, or proteins that change localization in sumoylation-defective cells (Cremona et al., 2012; Srikumar et al., 2013). We imaged each of these proteins tagged with GFP following 200 mM HU or 0.03% MMS treatment on an automated high-content screening confocal microscope system at seven times over four hours (Figure 2a). Having demonstrated the utility of an intensity distribution approach, we quantified protein re-localization in our HU and MMS screens. We identified 275 proteins that change subcellular location in HU and/ or MMS conditions (Figure 2b). We detected re-localization of 46 proteins for which replication stress induced relocalization had not been previously found in large-scale analyses, 25 of which were individually confirmed by visual inspection (Figure 2c and Table EV2). Of the remaining 21 proteins that did not visually validate, two proteins had low GFP fluorescence intensity, and 16 had high GFP intensity and showed a high degree of localization in untreated conditions (Table EV2). This suggests that these proteins may have displayed subtle changes in their intensity distribution not visible by eye yet were detected by our pipeline. We also improved the annotations for 62 proteins previously found to re-localize in MMS only that we find localize in both MMS and HU, and 38 proteins previously found to re-localize in HU only that we find localize in both HU and MMS. Thus, the core of proteins that respond to both types of DNA replication stress is more extensive than previously thought, and the degree to which the responses differ is less extensive, although as discussed below, large variations in the kinetics of response remain apparent, suggesting considerable stress-specificity of protein re-localization.

**Figure 2:**
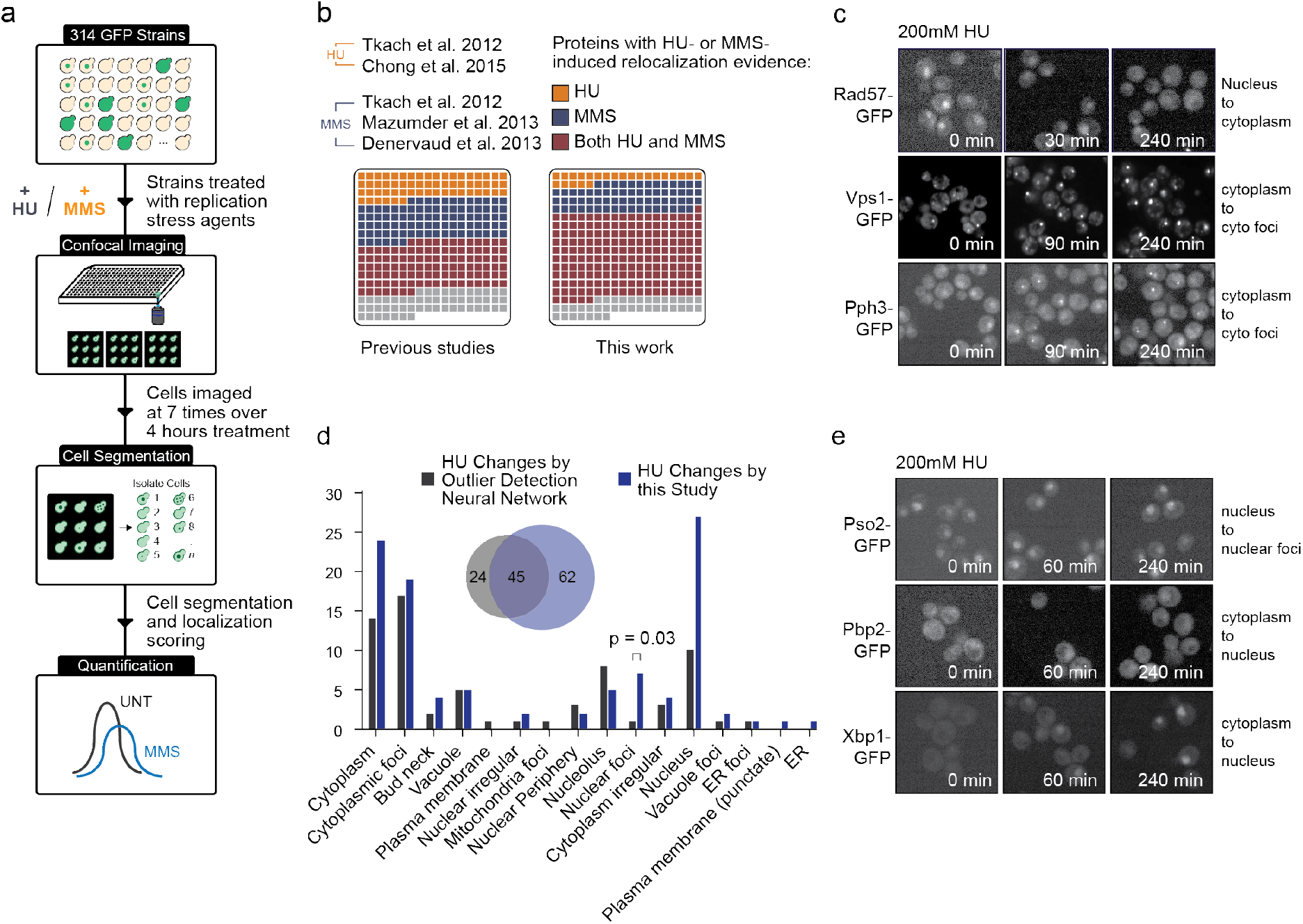
Quantifying pixel intensity distribution changes for 314 proteins uncovers new protein localizations in replication stress. (a) A set of 314 protein-GFP strains were imaged at 7 timepoints spanning 240 minutes following 0.03% MMS or 200 mM HU treatment. Cells were imaged on high throughput confocal microscope. Resulting images were subsequently segmented with CellProfiler (McQuin et al., 2018) and cells with protein re-localization events were measured with our quantification scheme. (b) The proteins assessed in our study are represented as individual squares. Those proteins identified to localize in HU only (orange), MMS only (blue), or both (red) from all the indicated literature data (left) and our imaging screen (right). (c) Representative micrographs of Rad57-, Vps1-, and Pph3-GFP re-localizing in 200 mM HU treatment. (d) Comparisons of the subcellular locations of each protein identified to change localization in machine learning analyses (Chong et al., grey) and our intensity distribution change method (blue). (e) Representative images of Pso2-, Pbp2-, and Xbp1-GFP protein re-localization after 200 mM HU treatment.

The four previous independent large-scale studies collectively detected HU- and/or MMS-induced localization changes in 260 of the 314 proteins in our study (Figure 2b) (Chong et al., 2015; Dénervaud et al., 2013; Mazumder et al., 2013; Tkach et al., 2012). Thirty of these proteins were not detected by our analysis. Of these proteins, 18/30 had high levels of localization in untreated conditions, which may result in the underestimation of the penetrance of protein localization following stress conditions as described above. Finally, the remaining nine proteins on our array were not detected in any analysis.

We asked if there were categories of localizations for which we were able to detect previously undescribed location changes, focusing on the 106 proteins that we found re-localized in HU. Computational analysis of localization changes in the yeast GFP collection following treatment with HU previously identified 69 proteins (Chong et al., 2015), and we detected 45 of these (a substantial overlap, hypergeometric p = 1.1 × 10-9). We detected more localization changes in proteins forming nuclear foci (Figure 2d), likely because the analysis used by Chong et al. did not include ‘nuclear foci’ as part of their neural network training. Perhaps more surprisingly, we also detected additional distributions to the cytoplasm (10 proteins) or nucleus (17 proteins). Conversely, and as noted above, Chong et al detected more localization changes for mitochondrial puncta proteins (Figure 2d), indicating the superior performance of outlier detection for this localization. Focussing on the proteins rather than the localization compartments, we identified 62 proteins that changed locations that were undetected in Chong et al. Visual inspection confirmed our image scoring in 44 cases (71%; Figure 2d and Table EV3). Interestingly, visual inspection of images for these 44 proteins downloaded from the CYCLoPS database (Koh et al., 2015) indicated that 31 show localization changes. We infer that the new HU-induced re-localizations that we detected are partially due to the stringent criteria for a localization change in Chong et al, leading to false negatives. We also note that Chong et al detected re-localization of 24 proteins that were negative in our screen (Table EV4). Visual inspection revealed that 16 of the proteins change localization in our images and 18 of these proteins change in images from Chong et al., indicating that 16 to 18 were false negative in our analysis. Five of these 18 proteins have complex localization patterns (nuclear periphery, endoplasmic reticulum, and mitochondria), which our pipeline scheme has difficulty quantifying, and 7/18 proteins have bright, intense GFP signal in untreated and treated conditions, where the pixel intensity distribution does not change, but the spatial patterning of the protein does, resulting in superior detection by machine learning methods. The differences between analysis methods reveal that gains can be realized by applying distinct analytical approaches to biological image data, and indicate that changes in study design can identify additional proteins that respond to replication stress, and uncover broad stress-induced localization changes for proteins that were previously detected in only one replication stress condition (Figure 2e).

### Diverse temporal responses to DNA replication stress

The time-scale of cellular responses to stress, including DNA replication, proteotoxic, and heat-induced stress, is well-characterized at the transcriptional level for cell populations (Dohn et al., 2021; Gasch et al., 2017; Rendleman et al., 2018). High-resolution analysis of the relocalization of 110 proteins in yeast revealed that the timing of re-localization events can be different in different stresses, can take place over extended periods of time, and can be transient (Dénervaud et al., 2013). These data indicate the utility of collecting data at multiple times to present a dynamic high-resolution view of protein location changes with high sensitivity.

Using our expanded dataset, spanning 212 relocalization events in HU and 256 in MMS, we asked if different kinetic re-localization patterns were evident. However, analysis of localization patterns is confounded by differences in the fraction of cells displaying protein re-localization in the population. We find considerable variation in the maximum percent of cells that display re-localization events (Figure 3a, Table EV5), that is apparent in either type of DNA replication stress. We applied a normalization to allow comparison of localization change kinetics independent of the fraction of cells displaying the localization change. The effect of the normalization can be seen in Figure 3b, where the maximum percent of cells with MMS-induced nuclear Ddc2 foci is 40.3% and the normalized % of maximum localization change is 100%, at t=180 minutes. After normalizing all the re-localization patterns, we used kmeans clustering to group proteins by similarity of relocalization pattern. The analysis converged on 5 groups of patterns for HU and 4 for MMS (Figure 3c, Table EV6), and thus the protein re-localization response does not occur with one temporal pattern. Although there are different re-localization patterns and they can be grouped based on pattern similarity, we were unable to detect any functional enrichments within any of the groups, including GO annotations, protein-protein interactions, genetic interactions, protein abundance, or intracellular compartment. We note that the re-localization kinetics of any protein is independent of the type of localization (for example, a protein moving into nuclear foci could do so with the same kinetics as a protein moving to the plasma membrane), and so it is not unexpected that functionally diverse proteins can share re-localization kinetic patterns.

**Figure 3:**
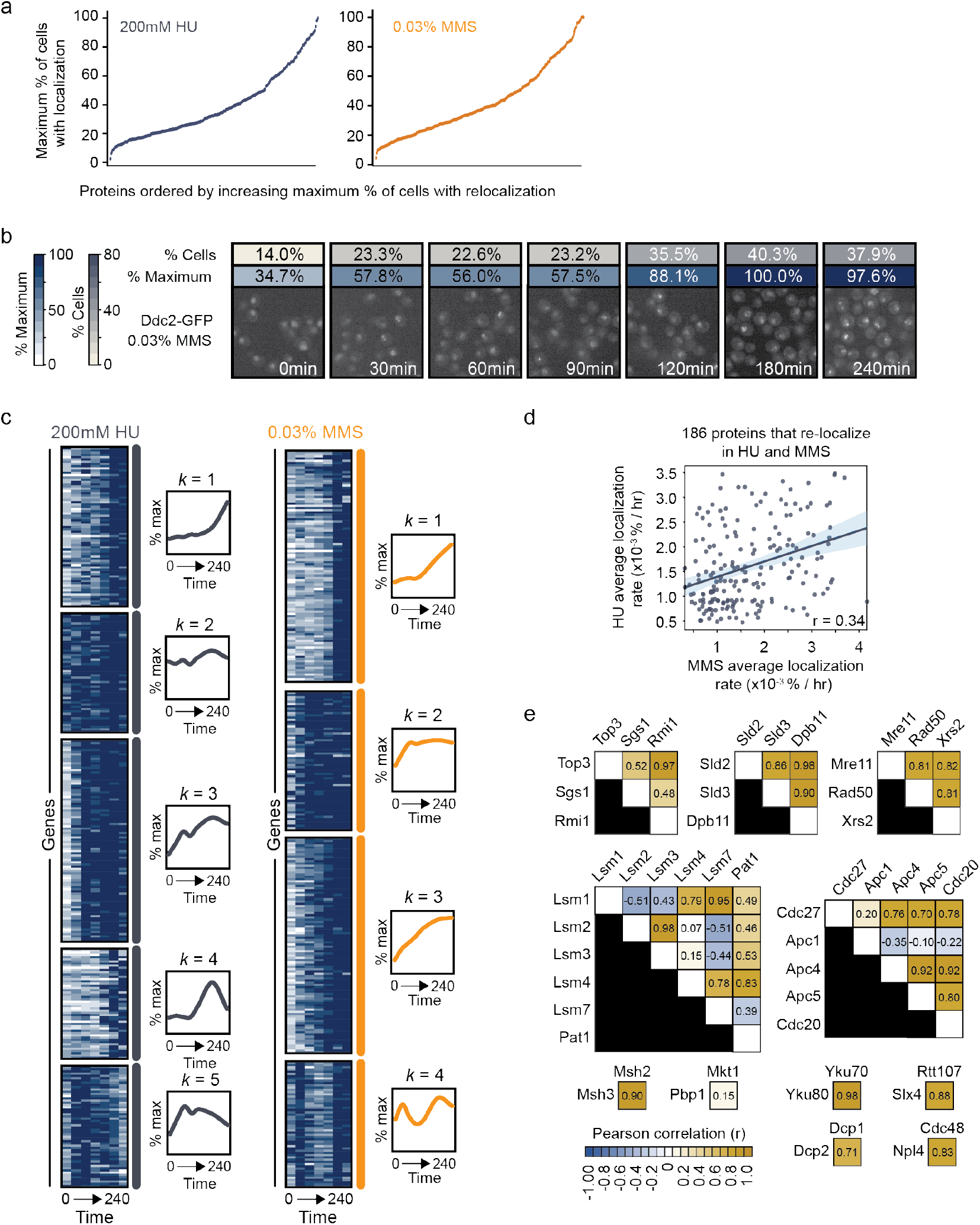
Replication stress induces diverse dynamics of protein re-localization. (a) The maximum penetrance, represented as the maximum % of cells with a localization change event at any timepoint in the drug treatment, is plotted for both 200 mM HU and 0.03% MMS conditions, and proteins are ordered by increasing maximum penetrance. (b) Representative images of the quantification pipeline applied to Ddc2-GFP focus formation after cells were treated with 0.03% MMS for 4 hours. Cells were imaged at 0, 30, 60, 90, 120, 180, and 240 minutes after MMS treatment. The degree of localization is indicated above the micrographs as the percentage of cells exhibiting a Ddc2-GFP focus (grey). To normalize dynamics of protein re-localization, the % of maximum was also calculated, where each value is divided by the maximum percentage of cells with a localization change (blue). (c) Heatmaps of the dynamics of protein localization in HU or MMS (calculated as % of maximum localization). Proteins are grouped by k-means clustering. Line plots illustrated to the right of each group of shows the average localization at each time point for all proteins within that k-cluster. (d) For the 186 proteins that were identified to re-localize in both MMS and HU conditions in our study, the average localization rate was calculated in each drug treatment and plotted against one another. The average re-localization rate was defined as the difference in the percent of cells with a change protein localization divided by 240 minutes of drug treatment. The shaded region represents the 95 percent confidence interval, and the Pearson correlation coefficient is displayed at the bottom of plot. (e) Correlation matrices for 11 protein complexes represented in our set of proteins analyzed as identified by Complex Portal, displaying the pairwise Pearson correlation coefficients for each protein member of the complex.

Analysing re-localization at multiple time points allowed us to identify localizations that were falsenegatives in previous analyses (Figure 2d, 2e). One effect of the improved annotation is that the apparent stressspecificity of protein re-localization is reduced: 186 of 314 proteins re-localize in both HU- and MMS-induced replication stress. We asked if stress-specificity was ev- ident in the re-localization kinetic patterns. When we compared the average re-localizaton rates of the set of proteins that re-localized in both HU and MMS conditions, we found that re-localization kinetics in MMS could only explain 34% of the re-localization kinetics in HU (Figure 3d). Furthermore, we found a substantial number of these proteins displayed poor correlation of localization rates between HU- and MMS-induced replication stress (69/186) proteins had a correlation coefficient less than 0.5, Figure EV2a). Along these lines, we calculated the fold-change differences between MMS and HU for each of these proteins across all time points and identify groups of proteins that indeed have different kinetic patterns between the two drug treatments (Figure EV2b, S2c), despite the proteins responding in both conditions Thus, re-localization kinetics are indeed stress-specific, even for related stresses like HU and MMS.

Despite the apparent absence of functional relationships in the kinetic pattern groups, we speculated that proteins that are co-complex members would show similar re-localization kinetics. We used Complex Portal (https://www.ebi.ac.uk/complexportal; accessed October 2021) to identify the functional protein complexes present in the set of 314 proteins that we analyzed. We then compared the re-localization kinetics of the members of the 11 protein complexes in MMS (Figure 3e) and in HU (Figure EV3a). We found that the pairwise re-localization kinetics correlated well among many complex members, and that their correlation was much higher than in protein pairs for same number complexes selected at random from the 314 proteins analyzed (Figure EV3b). However, we found interesting exceptions to the positive correlations of re-localization kinetics for cocomplex members. In MMS, the localization kinetics of Lsm2 and Lsm3 negatively correlate with those of Lsm1 and Lsm7 (Figure 3e). While this is surprising given that Lsm1-7 are thought to function as a heteroheptamer (Sharif and Conti, 2013; Wilusz and Wilusz, 2013) and that all localize to cytoplasmic P-bodies in diverse stresses (Decker and Parker, 2012; Dénervaud et al., 2013; Loll-Krippleber and Brown, 2017; Nissan and Parker, 2008; Teixeira and Parker, 2007; Tkach et al., 2012), it has been noted that their re-localization kinetics can differ (Dénervaud et al., 2013). Additionally, localization kinetics of Top3 more closely resembles those of Rmi1 than the kinetics of either Top3 or Rmi1 resembles those of co-complex member Sgs1 (Figure 3e). Interestingly, although Sgs1, Top3, and Rmi1 clearly function together in dissolution of recombination intermediates (Cejka et al., 2010; Wu et al., 2006; Yang et al., 2010), they have non-overlapping functions as *top3*Δ and *rmi1*Δstrains have substantial fitness defects that are absent in *sgs1*Δ (Chang et al., 2005; Gangloff et al., 1994). Finally, the localization trajectory of the anaphase promoting complex, Apc1, is negatively correlated with those for co-complex members Apc4 and Cdc20 (Figure 3e). We infer that differences in localization kinetics between accurately annotated complex members can reveal differences in protein function, and potentially underappreciate modes of complex regulation. Nonetheless, given that complex members tend to have similar re-localizations trajectories, we suggest that stratifying the re-localization kinetic data according to protein properties like complex membership could allow functional predictions to be made.

### Ydr132c/Mrx16 localizes to intranuclear quality (INQ) control foci in MMS

We reasoned that proteins with functional relationships should share re-localization kinetic trajectories and should re-localize to the same cellular compartment. We focused on MMS cluster 2, as it contains many proteins that localize to nuclear foci, a localization of particular interest in the DNA replication stress response (Figure 4a). From cluster 2, we took the subset of proteins that re-localize to nuclear foci, and then constructed a network where the nuclear foci proteins (the nodes in Figure 4a) are connected by edges that reflect the correlation of re-localization kinetics in both MMS and HU stresses. We noted that multiple components of the Intra-Nuclear Quality control (INQ) (Gallina et al., 2015; Mathew et al., 2020) body were highly correlated with a protein, Mrx16, that is annotated as associating with mitochondrial ribosomes (Möller-Hergt et al., 2018) yet localizes to the nucleus (Breker et al., 2014; Tkach et al., 2012). We tested Mrx16 co-localization with Nup49, Nop56, and Cmr1, members of the nuclear pore, nucleolus, and INQ body, respectively (Figure 4b). After treatment with MMS, Mrx16 clearly co-localized with Cmr1, and not Nop56 or Nup49. We quantified the co-localization, and found that Mrx16 foci form in 33% of cells, and 31% of Mrx16 foci colocalize with Cmr1 (Figure 4c). Cmr1 and other INQ proteins also form INQ bodies when cells are treated with the proteasome inhibitor MG-132 (Gallina et al., 2015). We found that Mrx16 responded similarly to Cmr1, forming foci following treatment with MG-132 (Figure 4d), further indicating that Mrx16 foci correspond to INQ bodies. We infer that Mrx16 is a previously uncharacterized component of INQ bodies, and that protein re-localization kinetics patterns can be stratified to predict protein function.

**Figure 4:**
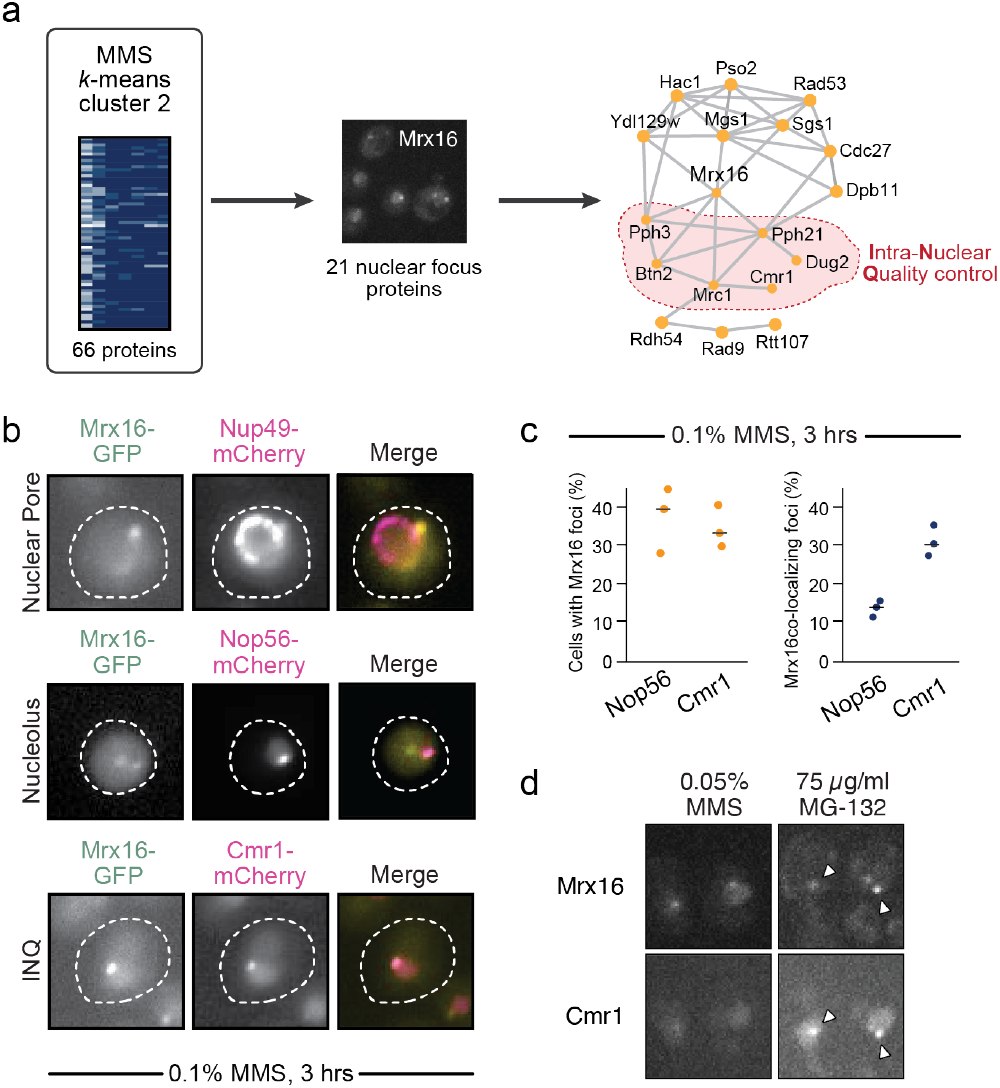
Previously undetected Mrx16 forms nuclear foci that co-localizes with the INQ compartment. (a) From the 66 proteins in cluster k-2 in MMS treatment, 21 proteins form nuclear foci in replication stress, and these Pearson correlation coefficients (r) for these were plotted as a network. Nodes represent proteins, edges connect proteins with the most similar r values, and the red region highlights proteins previously associated with the INQ compartment. (b) Representative micrographs of Mrx16-GFP in yellow after treatment of cells with 0.1% MMS for 3 hours. Shown in magenta is Nup49-, Nop56-, and Cmr1mCherry to mark the nuclear pore, nucleolus, and INQ compartments, respectively. (c) Micrograph images of Mrx16-GFP and Cmr1-mCherry, with arrows indicating the location of each respective nuclear focus. A merged image is displayed, with the white arrows indicating the location of Mrx16-GFP and Cmr1-mCherry co-localized foci. The quantification for proportion of cells that exhibit Ydr132c/Mrx16-GFP focus formation and co-localized foci with the respective marker are displayed below. (d) Confocal micrographs of nuclear focus formation of the indicated proteins during 0.05% MMS or 75ug/mL MG-132 treatment.

### Heterogeneity of protein re-localization in response to stress

One product of our analysis is a cell-by-cell evaluation of localization change for each protein, allowing us to assess penetrance (the proportion of an isogenic population that expresses the given phenotype) for 468 protein re-localization phenotypes related to DNA replication stress response. As indicated in Figure 3a, we observed a wide range of degrees of penetrance, where two genetically identical cells did not display the same protein localization, from a minimum of 14.1% of cells (Msh3 in HU) to a maximum of almost 100% (Hsp42 in HU and Ylr297w in MMS). Strikingly, we even note several proteins with extensive localization penetrance, ranging from 0-34.5% in unperturbed conditions (Figure 3a, 0 minutes MMS or HU treatment). We plotted the penetrance distributions (Figure 5a) and found that in either replication stress, the distribution is left-skewed with peaks at a relatively low penetrance of 20 – 30%. When comparing the two replication stresses, a positive trend is evident. However, there are numerous examples of protein re-localizations that show high penetrance in one stress and not in the other. It is also evident that there was not a systematic shift from one stress to the other that might reflect different stress levels imposed by the two agents at the concentrations used.

**Figure 5:**
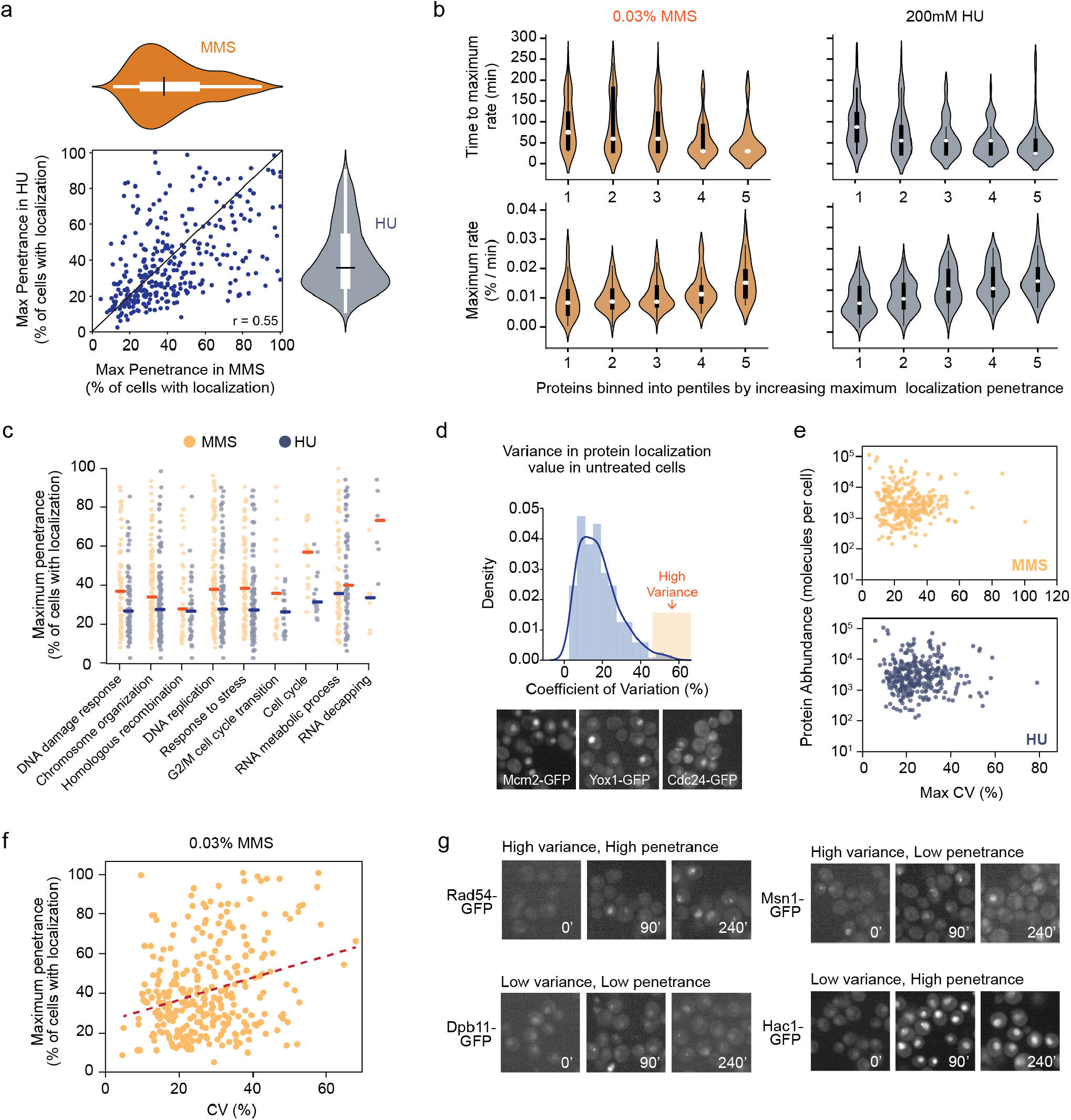
High degree of protein re-localization heterogeneity during replication stress. (a) The maximum penetrance (calculated as % of maximum localization change) for both HU and MMS are shown in the scatterplot. The distributions for maximum penetrance in each condition is shown on the respective plot axes as violin plots. The black line represents the median, and the large white rectangle represents the IQR (25th – 75th percentile). (b) Proteins were ordered by increasing maximum penetrance and binned into pentiles, where bin 1 represents 64 proteins with the lower penetrance, and bin 5 represents the highest penetrant proteins. Protein localization rates were calculated for each time point in the study, with the maximum rate of localization, and the time at which the maximum rate is exhibited, is plotted for each bin. (c) The maximum penetrance of protein re-localization was plotted for proteins in each GO biological process indicated, for both MMS (orange) and HU (blue). (d) The coefficient of variation (CV) was calculated for each cellular LOC score for each protein during unperturbed conditions. Proteins that exhibit high CV, and thus high variation in localization responses, is indicated in orange region on the right of the histogram (CV > 40%). Representative images of those proteins (Mcm2-, Yox1-, and Cdc24-GFP) are shown below. (e) Scatterplot of maximum CV for each protein compared to protein abundance levels, as calculated in Ho et al, 2018. (f) The maximum penetrance of protein re-localization is plotted versus the maximum coefficient of variation for each protein in MMS conditions. (h) Representative images of proteins that exhibit varying degrees of variance and penetrance. High penetrance (% max > 50), low penetrance (% max < 50), high variance (CV > 40%) and low variance (CV < 10%).

The relationship between phenotypic penetrance and kinetics has not been explored extensively. In amyloid diseases, protein unfolding kinetics contribute to penetrance, with faster unfolding correlating with more penetrant disease (Hammarström et al., 2002). How the timing of a phenotype might relate to phenotypic penetrance in other settings is unclear. We grouped proteins into 5 bins according to maximum penetrance during our imaging time courses. The rates of protein localization and the timepoints at which the maximum rate was observed were then plotted for each bin to reveal the average kinetics of re-localization (Figure 5b). We found that the fastest responses tended to be the most penetrant, in either MMS or HU stress. Conversely, less penetrant responses tended to be slower.

We next asked if proteins with related functions displayed similar penetrance in HU and MMS, examining GO biological processes related to the DNA replication stress response (Figure 5c). In most cases, the median maximum penetrance was similar in HU and MMS. Two notable exceptions emerged. Proteins annotated to ‘G2/M cell cycle transition’ had higher phenotypic penetrance in MMS than in HU, likely because HU stress poses a stronger barrier to S phase progression than does MMS, and MMS treated cells accumulate at G2/M due to replication checkpoint activation (Kupiec and Simchen, 1985; O’Connell et al., 2000; Pellicioli et al., 2001; Yamamoto et al., 1996). The second exception is ‘RNA decapping’ proteins, which have much higher penetrance in HU than they do in MMS. While the physiological basis for this difference is unknown, we and others have noted that cytoplasmic foci referred to as P-bodies (the relevant localization for decapping proteins in stressed cells) form more rapidly and readily in HU than in MMS (Dénervaud et al., 2013; Tkach et al., 2012). In summary, we find extensive heterogeneity in the extent of phenotypic penetrance for different protein re-localizations, and there appears to be a high degree of stress-specificity associated with phenotypic penetrance when considering protein localization changes.

When we previously examined phenotypic penetrance, we reduced the calculated LOC score for the cellular phenotype (protein re-localization) of a given protein to two categories: re-localized or not. However, since several localization parameters can influence the LOC value, including signal intensity and size, proteins within a cell population that localize to the same compartment may present different LOC values. We reasoned that measuring the variation of the LOC value itself would provide insight into the heterogeneity of the localization response. Therefore, we measured the coefficient of variation (CV) of the LOC values for each protein at each timepoint. As previously noted, unperturbed cells displayed phenotypic heterogeneity. We therefore first examined the distribution of the CV for unperturbed cells (Figure 5d, Table EV7). A group of 13 proteins with very high variance (CV>40%) was apparent. Inspection of the relevant images revealed that in each case a large fraction of the cells were expressing the re-localized phenotype in the absence of perturbation. Therefore, some re-localization phenotypes are characterized by very strong prior variation in the untreated cells, reminiscent of ‘bet-hedging’ strategies where a fraction of a cell population pre-expresses a potentially desirable state. The high-variance-in-untreated group includes proteins whose localization is cell cycle regulated (Figure 5d; Mcm2, Yox1, Cdc24). Another protein in the highvariance-in-untreated group, Msn2, is known to show pulsatile behaviour (Dalal et al., 2014). The basis for extensive re-localization in the absence of perturbation for the remaining nine proteins remains to be defined.

Heterogeneity in gene expression often underlies phenotypic heterogeneity (Levy et al., 2012; Liu et al., 2019; Newman et al., 2006) and is, in part, attributed to fluctuations in the molecular compositions of individual cells due to stochastic partitioning of molecules between daughter cells. Since heterogeneity in such a model would be favoured by low protein abundance, we asked if there was a relationship between protein localization heterogeneity and protein abundance. Correlation plots of the maximum CV for each protein versus the abundance of each protein showed a substantial range of abundances for proteins with the most heterogeneous localization patterns (Figure 5e). We infer that heterogeneity in stress-induced protein re-localization is not principally driven by low protein abundance.

Finally, we measured the CV for the LOC scores for each protein during MMS stress and compared this measurement of heterogeneity to phenotypic penetrance (fraction of cells showing localization change) (Figure 5f). There is a general correlation between variation in localization and the extent of penetrance, suggesting that the most penetrant protein re-localization responses reflect localization states that are themselves highly variable. An example of high variation/high penetrance is Rad54 (Figure 5g) where most cells display re-localization, but the intensity and number of the protein locations (in this case nuclear foci) is highly variable. The reciprocal case, where both variation and penetrance are low reflects proteins where the fraction of cells that display re-localization is small, and the fluorescence intensity of the relocalization compartment is low, for example Dpb11 (Figure 5g). Nonetheless, we find that variation can be independent of penetrance (Figure 5g). The low variation group contains proteins with highly penetrant localization changes, for example Hac1, which re-localizes very uniformly to the nucleus. The high variation group contains proteins with low re-localization penetrance, like Msn1, where localization to the nucleus is highly variable from cell to cell and occurs in only small subset of cells. Overall, cells respond to replication stress with high variability at the level of proteome re-localization. The variable responses in subsets of the population and their influence on other phenotypes (e.g., cellular fitness to stress) for many of these proteins will be critical for a complete understanding of the cellular response to stress.

### Population localization dynamics reflect the average of all component single-cell localization trajectories

Thus far, our measurements of cellular heterogeneity have been based on snap shots of different single cells within a population before and after replication stress conditions. This allowed us to determine the presence and magnitude of heterogeneity in a population of cells but failed to capture heterogeneity at the individual cell level over time. To this end, we measured the localization response of the same single cells over time in MMS conditions with greater time resolution by acquiring images every 7.5 minutes, and tracking each individual cell during the course of the experiment. Specifically, we collected time-lapse images of yeast cells in stationary phase grown in a CellClamper microfluidic system (Schmidt et al., 2018) for four hours unperturbed (acclimatization period) and then an additional four hours following MMS treatment. We chose to monitor Lsm1 and Lsm7 cytoplasmic focus formation since they showed high correlation in MMS (Figure 3e) and are part of the same macromolecular complex. We calculated the percent of cells within the population displaying Lsm7 foci using the same intensity distribution scheme described above. We found that Lsm7 makes foci in stationary phase, which eventually dissolve as the cells resume logarithmic growth, and then quickly re-form once again following the addition of MMS (Figure 6a). Since our microfluidic imaging platform permits single cell tracking over time, we can deconstruct population-based localization quantifications into singlecell components (see methods), following an individual cell’s protein re-localization response throughout the entire experiment (Figure 6b). It is immediately clear that there is substantial variation in cell localization dynamics. The time when Lsm7 re-localizes to foci is drastically different between cells, with some cells displaying almost instantaneous re-localization and some being delayed for up to 175 minutes. Of the 50 cells examined, 2 do not show any re-localization following MMS treatment, and 2 cells appear to show constant Lsm7 foci independent of treatment (Figure 6b). Similar observations were made with Lsm1 localization (Figure 6c, 6d), however an important difference noted was that Lsm7 shows a greater proportion of cells with a phenotype in untreated conditions, illustrating how heterogeneity can be protein specific even for members of the same complex. Perhaps more interesting was the presence of a small population of cells, for both Lsm1 and Lsm7, that did not show any evidence of localization throughout the entire treatment. The reason for this lack of response to replication stress is unclear, but the fitness and other cellular phenotypes of these cells is certainly of interest. Overall, we find that the extensive phenotypic variation we observe within the cellular population is the result of high variation among its single cell components.

**Figure 6:**
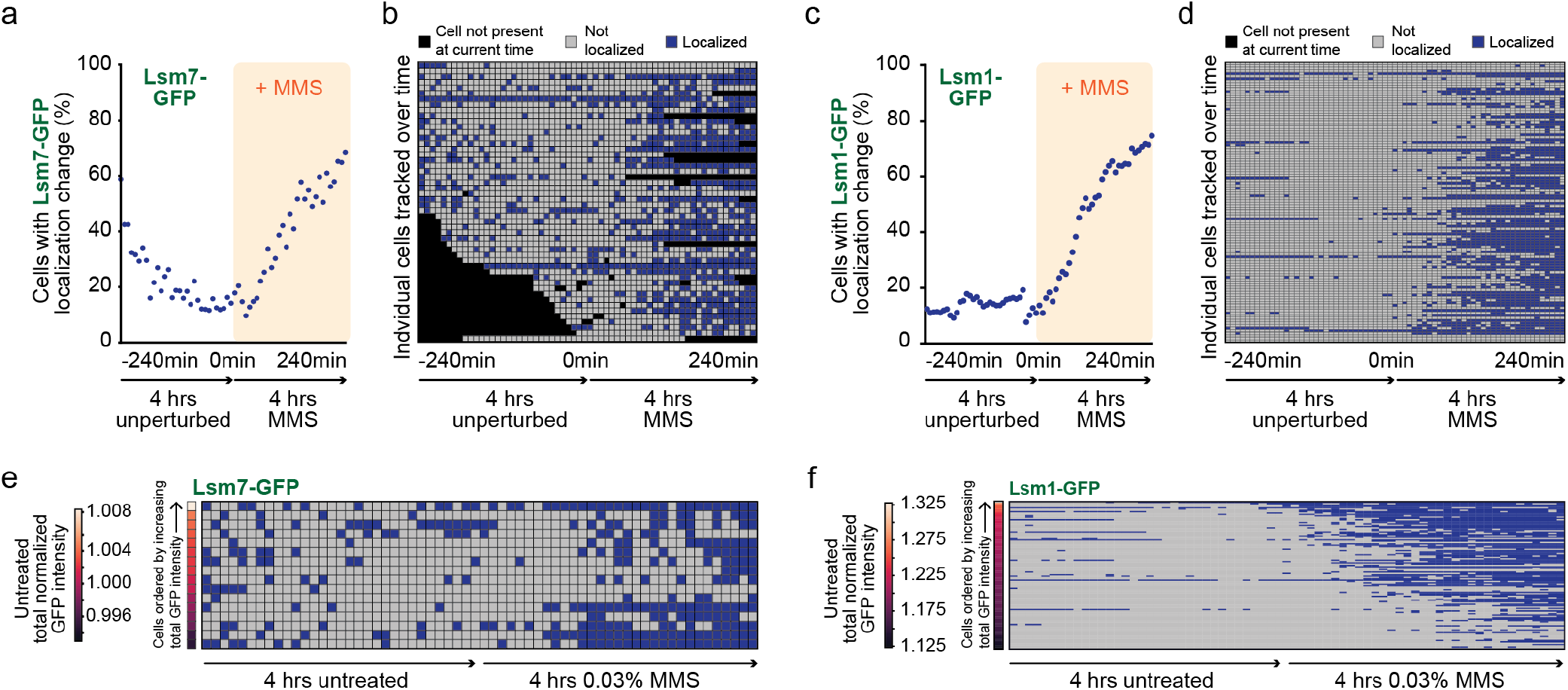
Factors influencing the extensive phenotypic heterogeneity during replication stress at the single cell level is protein specific. (a) Cells were imaged in the CellClamper microfluidic device (Schmidt et al., 2018). Cells were added to the device with a constant supply of YPD media supplied. Cells were imaged on an immunofluorescence microscope for 8 hours, with images acquired every 7.5 minutes,for 4 hours in YPD, and 4 hours in YPD plus 0.03% MMS. The % of cells with Lsm7-GFP cytoplasmic foci was calculated and plotted over time. (b) Heatmap showing the trajectory of each individual cell in the population of Lsm7-GFP cells that was quantified in (a). Black squares represent cells that have not been detected (have not yet divided from the mother cell) at that point in the experiment. Grey squares are cells that have not shown a Lsm7-GFP localization change, and blue squares are cells that are changed in Lsm7-GFP localization. (c) The % of cells and (d) the single-cell re-localization measurements for Lsm1-GFP were quantified in the same manner. (e) Cells that persisted throughout the entire 8-hour time course in the microfluidic system are shown as a heatmap. The mean total normalized Lsm7-GFP intensity was calculated for each cell during unperturbed conditions, and the heatmap is sorted by increasing GFP intensity. (f) This was similarly performed for Lsm1-GFP.

The relation between gene expression, protein abundance, and phenotype differences have been observed before (Liu et al., 2019; Uphoff et al., 2016; Vincent and Uphoff, 2021) but the relation between protein expression level and protein subcellular localization dynamics is unclear. We asked if protein abundance could drive re-localization by measuring the abundance of Lsm7 and Lsm1 (total GFP intensity) in untreated conditions for each individual cell tracked over the entire time course, and related the protein abundance to protein localization kinetics. With Lsm7-GFP, there is no clear relationship between protein abundance and a cell’s re-localization dynamics (Figure 6e). This is in stark contrast to Lsm1-GFP, where cells with lower levels of Lsm1-GFP protein form cytoplasmic foci even at later times, whereas those cells with higher abundance make foci more quickly following stress (Figure 6f). Therefore, while protein abundance can contribute to protein localization phenotypes, we provide evidence that this can be protein specific, even for functionally related proteins complex members.

## Discussion

The ability to measure subcellular proteomic changes has become increasingly important for a complete understanding of the cellular response to stress. Several studies have advanced image analysis pipelines to determine where proteins reside inside yeast cells (Almagro Armenteros et al., 2017; Chen et al., 2007b; Chong et al., 2015; Grys et al., 2017; Kraus et al., 2017; Lu et al., 2018; Ouyang et al., 2019; Xiao et al., 2019). However, these techniques rely upon additional data sets, either to train machine learning models (Chong et al., 2015) or for systematic comparisons of proteins to explore localization similarity among data sets (Lu et al., 2018). While powerful, the applicability of these methods requires additional image information that might not be readily available. Here, we have designed a method that accurately quantifies protein-GFP localization changes in populations of cells compared to a control population and have applied this quantification pipeline to cells exposed to replication stress induced by MMS or HU. Although our quantification requires prior knowledge of the involved cellular compartments, it efficiently captures changes in proteome spatial dynamics. Impressively, our pixel intensity distribution change detection is suitable for use on images acquired from a variety sources. We demonstrated that our method detects protein localization changes in images acquired from similar high-throughput confocal platforms, and from drastically different widefield microscopic images. Our quantification method is a powerful tool that we used to study the temporal changes in the yeast proteome following DNA replication stress.

### Replication stress results in diverse protein relocalization dynamics

Previous studies of protein localization during replication stress conditions have provided a comprehensive view of the proteins that respond to replication stress (Chong et al., 2015; Dénervaud et al., 2013; Mazumder et al., 2013; Tkach et al., 2012). With this defined set of proteins, we performed high throughput confocal imaging and applied our computational quantification to 314 proteins. One key advantage of our study is that we compared protein re-localization events in HU and MMS, which both cause replication stress by different mechanisms, providing the opportunity to examine drug-specific responses in protein localization dynamics. We followed movement of 314 proteins and identified localization changes that had not been detected before. We find that a greater proportion of proteins (68%) re-localize in both HU and MMS than was previously thought (Tkach et al., 2012). However, when the kinetic patterns of localization change are discerned, considerable stress specificity is observed even for highly related stresses like HU and MMS. Thus, we present a concrete illustration of the importance of measuring stress responses over multiple treatment times.

Moreover, we demonstrate that temporal changes in protein re-localization are an additional property that can be leveraged to infer protein function. We explored the function Mrx16, which was found associated with mitochondrial organization of gene expression (MIOREX) complexes through the purification of yeast mitochondria from cell lysates (Möller-Hergt et al., 2018). MIOREX complexes are evenly distributed across the entirety of the mitochondria (Möller-Hergt et al., 2018) and play a key role in spatially organizing the transcription, mRNA maturation, translation, and mRNA decay machinery of the mitochondria (Kehrein et al., 2015a, 2015b). Counter to expectations, when we combined quantified re-localization temporal patterns with cell compartment annotations, we found that Mrx16 is a component of the intranuclear quality control (INQ) compartment (Figure 4). Mrx16 formed nuclear foci upon MMS-induced replication stress with kinetics similar to those of known INQ proteins. We found that Mrx16 nuclear foci correspond to the INQ compartment, as they colocalize with a key INQ protein, Cmr1 (Gallina et al., 2015), and suggest that Mrx16 is a component of INQ, rather than a substrate.

### Phenotypic heterogeneity contributes to the replication stress response

It is widely accepted fact that genetically identical cells grown in the same culture exhibit remarkable cell-to-cell phenotypic differences. Cellular heterogeneity has been well documented at the level of gene expression and protein abundance, and a few genetic determinants have been identified in budding yeast. One of the first global analyses of protein abundance variation in an unperturbed population of yeast cells identified functional enrichment among proteins whose abundance exhibited high or low variation (Newman et al., 2006). Proteins that were generally part of stress responses or have chromatin related functions (i.e. chromatin remodelling and transcription factor binding) had large variations in abundance measurements. Phenotypic diversity can have important functional consequences, particularly during stress conditions where subpopulations of cells may be better poised to respond to stress. Our data suggest that phenotypic diversity extends beyond gene expression and protein abundance and presents itself at the level of protein subcellular localization as well.

Extensive phenotypic heterogeneity was observed for protein localization in response to HUand MMSinduced replication stress; numerous proteins had different localization penetrance between the two drug treatments. We showed that the response of a cell population to replication stress was the summation of all its component single-cell localization trajectories and, using Lsm1 and Lsm7 as examples, we demonstrated that individual single cells can exhibit high variability in localization dynamics (Figure 6b, d). Noteworthy was the observation that protein abundance influenced localization dynamics of Lsm1, but not of Lsm7, despite these two proteins being members of the exact same protein complex (Figure 6e, f). It has been shown previously that homologous recombination repair phenotypes seem to correlate with the expression level of genes that influence homologous recombination directly (measured by Rad52) or indirectly (measured with Rad27) (Liu et al., 2019). This may be true for several proteins, but as with the case for Lsm1 and Lsm7, the factors that influence cellular heterogeneity appear to be protein specific, given that protein abundance did not influence the localization dynamics of replication stress response proteins uniformly (Figures 5e, 6e, 6f).

In summary, we provide a method to capture changes in protein subcellular location following stress conditions, quantifying protein localization dynamics, and revealing the large phenotypic heterogeneity that is displayed during replication stress. Phenotypic heterogeneity is evident throughout evolution, but has been studied mainly at the level of mRNA expression and protein abundance. We extend the phenotypic heterogeneity space to include the complex molecular process of protein location and compartmentalization. Our work provides a framework to further explore the phenomena that contribute to protein localization heterogeneity, and to define the biological role of localization heterogeneity in cellular responses to stress.

## Methods Details

### Yeast Strains and Media

The 314 yeast strains with protein-GFP fusions used for the high-throughput confocal microscopic screen were derived from the yeast GFP collection. A SML1 null mutation was introduced into each of these GFP-tagged strains by performing Synthetic Genetic Array (SGA) with BHY021 as the query strain. These resulting strains were used for all high-throughput confocal microscopic screens. Yeast strains for all other experiments in this study are derivatives of BY4741. Standard yeast media and growth conditions were used for all experiments, unless indicated otherwise. Strains were constructed through genetic crosses and standard PCR-based disruption transformations.

### High-Throughput (HTP) Confocal Microscopy

The 322 GFP-tagged strains were grown to midlogarithmic phase in liquid low fluorescence media (LFM) (1.7g/L yeast nitrogenous base without ammonium sulfate, 1g/L monosodium glutamate, 0.02g/L uracil, 0.02g/L histidine, 0.1g/L leucine, 0.15g/L methionine) and imaged on a 384-well imaging plate (PerkinElmer 384 Cell Carrier Ultra, 6057300) in a high-throughput confocal microscope (PerkinElmer Opera High-Content Screening System) as previously described (Torres & Brown, 2015). Briefly, logarithmically-growing cells were added to the imaging plate to a density of OD600 = 0.02 and incubated at 30°C for 1 hour, without agitation, to allow cells to settle to the bottom of the wells. Cells were then treated with 200mM HU or 0.03% MMS and imaged at 7 timepoints over 4 hours (0, 30, 60, 90, 120, 180, and 240 minutes). The addition of drug to each column of the plate was staggered by 1 minute to compensate for the imaging time of the Opera microscope. For each time point, 4 fields in different regions of the well were imaged. Both GFP and RFP was imaged on the microscope with 800ms exposure.

### Protein Localization Quantification from HTP Confocal Microscopy

Cells expressed a constitutively cytoplasmic tdTomato fluorophore, and whole cell image segmentation was conducted using CellProfiler based on tdTomato signal. Dead cells displaying high GFP fluorescence were removed from the analysis. An automated quantification scheme was developed using custom written Python scripts to acquire measurements of GFP pixel intensities for each individual cell from all obtained images. The pixel intensity distributions for each cell were normalized by the median GFP intensity measured for each respective cell. First, for a given protein, the 95th percentile was determined from the pixel intensity distribution for each single cell (hereafter referred to as the localization, or LOC, value) in the untreated condition. The median (x) and median absolute distribution (MAD) was calculated for the LOC values among only the untreated population of cells for a given protein. These parameters were used to determine the threshold values for changes in protein localization. If a cell LOC value was greater than 2 MAD from the median (x + 2 MAD), the cell was considered to exhibit an increase in protein re-localization from the untreated condition. Conversely, if a cellular LOC value was less than 2 MAD from the median (x - 2 MAD), it was labeled as decreased localization from the untreated condition. Next, the LOC value was calculated for all cells among all proteins and across all 7 timepoints in our study. The percent of cells with a localization change compared to the untreated population of cells for the respective protein (either an increase or decrease) was calculated. Finally, we calculated a percent of maximum value (% max) to normalize the localization dynamics between proteins, which was the percent of cells at any timepoint divided by the maximum percentage at any given time point for that protein.

### Clustering Analysis

Clustering by k-means was performed for both calculations of the percent of re-localization and the % max values, with the indicated number of k clusters. To determine an appropriate number of k clusters, the total within-cluster sum of squares was measured with increasing numbers of clusters. The number of clusters were determined suitable for k-means clustering analysis since increasing the number of clusters did not provide better modelling of the data, as determined by the elbow test (measuring total within variability), gap test, and cluster tree test.

### Data Literature Data Collection and Comparisons Between Studies

We gathered three large data sets from published studies cataloguing global protein localization changes in stress conditions (Chong et al, 2015; Dénervaud et al, 2013; Breker et al, 2013). For analyses and visualization of the intersection of hits between these localization studies, only proteins in the literature studies that were also in our mini array of 314 genes were considered. Raw images from two of these studies (Chong et al, 2015; Dénervaud et al, 2013) were downloaded from their respective databases.

### Gene Ontology (GO) Term and GeneMANIA Interaction Enrichment Analysis

GO Term Finder version 0.86 (https://www.yeastgenome.org/goTermFinder) was used to identify enriched function terms (default settings p-value < 0.1 with false discovery rate being calculated). The background set of genes was restricted the 314 genes in our mini-array. The final background set consisted of 314 proteins.

### Nuclear Ydr132c/Mrx16 Focus Formation Microscopy

For imaging performed on the Nikon Eclipse Ti-2 Inverted Microscope: cells expressing Ydr132c/Mrx16-GFP (in the indicated genetic backgrounds) were grown in liquid YPD at 30°C to mid-logarithmic phase (OD600 = 0.3 – 0.6). A 1mL aliquot of untreated cells was taken for imaging. The remainder of the culture was treated with 0.03% MMS for 180 min, after which another 1mL aliquot was taken for imaging the MMS-treated cells. Each aliquot of cells was centrifuged at 4000 rpm for 2 minutes. Cells were resuspended in 1mL SD-All media (6.7g/L yeast nitrogenous base with ammonium sulfate, 1X amino acids, 2% w/v glucose). Cells were centrifuged again, resuspended with 500uL SD-All, and 1.5uL was spotted onto a glass slide with a coverslip and imaged on the microscope using Micro-Manager Imaging Software. Images in GFP, RFP, and DIC channels were acquired for Ydr132c/Mrx16-GFP, protein-mCherry fusions, and cell morphology, respectively. The resulting images were visually inspected and quantified by counting number of cells with at least one Ydr132c/Mrx16-GFP focus.

For imaging performed in the PerkinElmer Opera Phenix: cells were grown in liquid SD-All at 30°C to mid-logarithmic phase (OD600 = 0.3 – 0.6), and 0.05 ODs were added to a 384-well imaging plate to a final volume of 90uL and incubated at 30°C to allow cells to settle to the bottom of the plate. After 1h, 10uL of 10X MMS or HU was added to the wells, and the plate was imaged for 4 hours every 15 minutes.

### Microfluidic Time-Lapse Imaging and Analysis

We used the CellClamper Microfluidic Device for timelapse fluorescence imaging of single cells tracked over time (Schmidt et al., 2018). The cleaning and set up of the CellClamper was performed as previously described. Cells were grown in liquid YPD at 30°C overnight, diluted to 5.0 × 108 cells/mL, and 0.67uL was added to the PDMS culture pad of the device. Liquid YPD media was delivered from a reservoir to cells by gravity flow. The cells were allowed to acclimatize and grow on the device for 4 hours, after which the reservoir was replaced with YPD + 0.03% MMS. Cells were allowed to grow in MMS for an additional 4 hours. Throughout the 8 hour time course, images were collected every 7.5 minutes using GFP, RFP, and cell differential interference contrast filter sets (Thor labs). Cell segmentation and tracking was performed using CellX software as previously described (Mayer et al., 2013). The resulting image masks outputted from CellX was used to identify the single cells that were tracked throughout the entire experiment. Using the same pixel intensity quantification scheme as described above, LOC values were calculated for each cell. The parameters for the CellX segmentation are provided on the online GitHub repository.

### Quantification and Statistical Analysis

All statistical analysis, data manipulation, and data visualization was performed in Python (https://www.python.org/) and R (https://www.rproject.org). All the details of data analysis can be found in the Results and Materials and Method sections.

### Data and Software Availability

Datasets are provided in the supplemental tables S1 through S7. The source code supporting the conclusions and data visualization of this article is available in the GitHub repository, at https://github.com/bqho/localizationQuantification. Complete raw image data available from the authors upon request.

## Acknowledgements

This work was supported by the Canadian Institutes for Health Research (FDN-159913 to GWB), an Ontario Government Scholarship and a Natural Sciences and Engineering Research Council of Canada CGS-D award (to B.H.). GWB holds a Canada Research Chair (Tier 1). We thank Yoshikazu Ohya, Tajinder Ubhi, Peter Bartlett, Charles Boone, and Brenda Andrews for helpful discussions and careful reading of the manuscript. We are grateful to work on the lands of the Mississaugas of the Credit, the Anishnaabeg, the Haudenosaunee and the Wendat peoples, land that is now home to many diverse First Nations, Inuit, and Métis peoples.

## Author Contributions

Conceptualization, B.H., N.P.T., and G.W.B; Methodology, B.H., N.P.T., R.L.K., and G.W.B; Microscopy Experiments, B.H., N.P.T., A.C; Formal Analysis, B.H., A.C., F.R., and G.W.B.; Writing, B.H. and G.W.B.; Writing – Review & Editing, B.H., R.L.K., A.C., F.R., and G.W.B.

---

**Figure S1:**
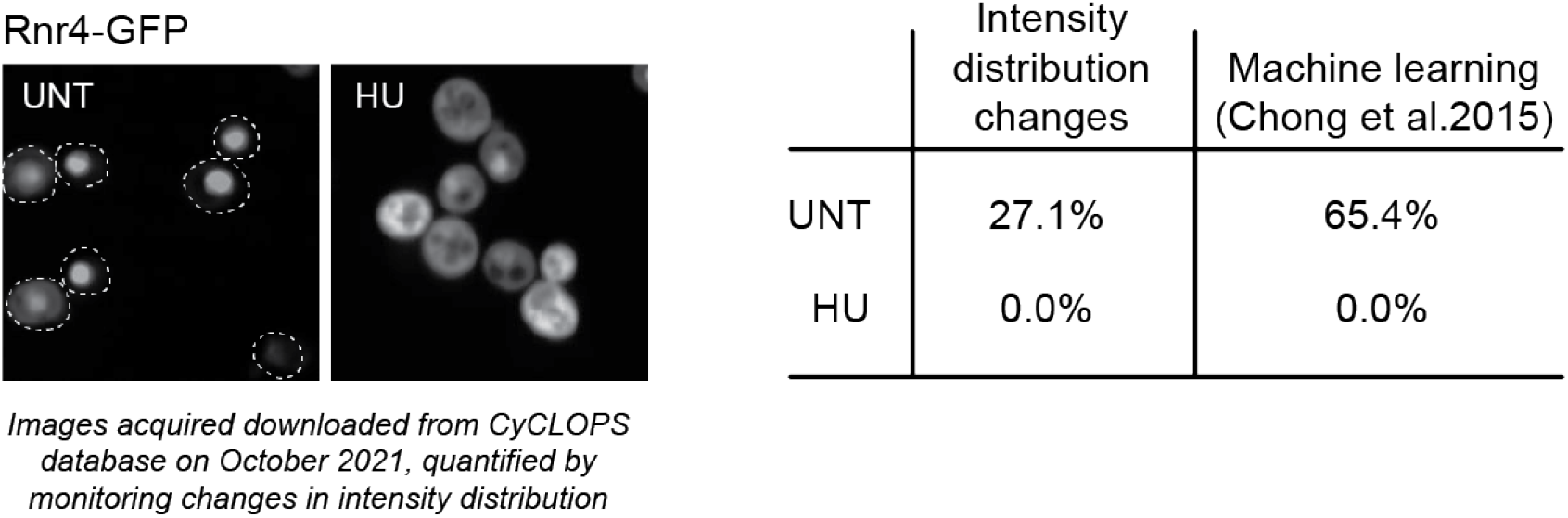
Intensity distribution changes can underestimate protein re-localization. Representative images of Rnr4-GFP re-localization from the nucleus (in untreated, UNT) to the cytoplasm (in HU) were acquired from the CYCLoPs database and are shown. White dotted outlines represent the whole cell. These images were scored by my intensity distribution change pipeline for the percent of cells with Rnr4-GFP in the nucleus, and were compared to the values calculated by Chong et al.

**Figure S2:**
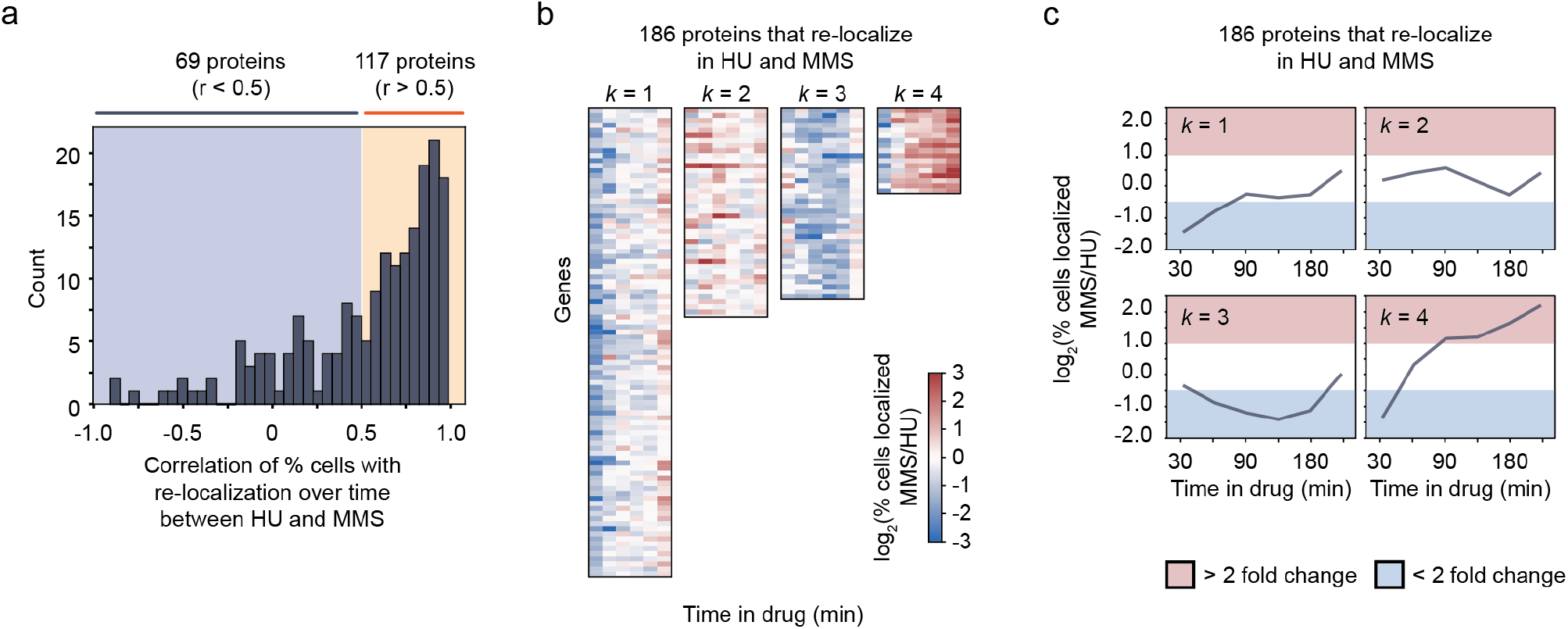
Proteins that re-localize in MMS and HU conditions exhibit stress-specific kinetics. (a) The Pearson correlation coefficient of the localization changes over time for each of the 186 proteins that re-localize in both MMS and HU conditions is plotted as a histogram. The blue and red shaded regions correspond to proteins with low (r < 0.5) and high (r>0.5) correlation coefficients. (b) The log2(fold-change) between the percent of cells with re-localization in MMS vs. HU conditions was calculated for the 186 proteins at which time point, and clustered into 4 groups by k-means algorithm. (c) The average log2(fold-change) among the proteins within each of these k-groups is plotted over time. Red and blue shaded regions of the plots represent greater or less than two-fold change in the percent of cells with re-localization.

**Figure S3:**
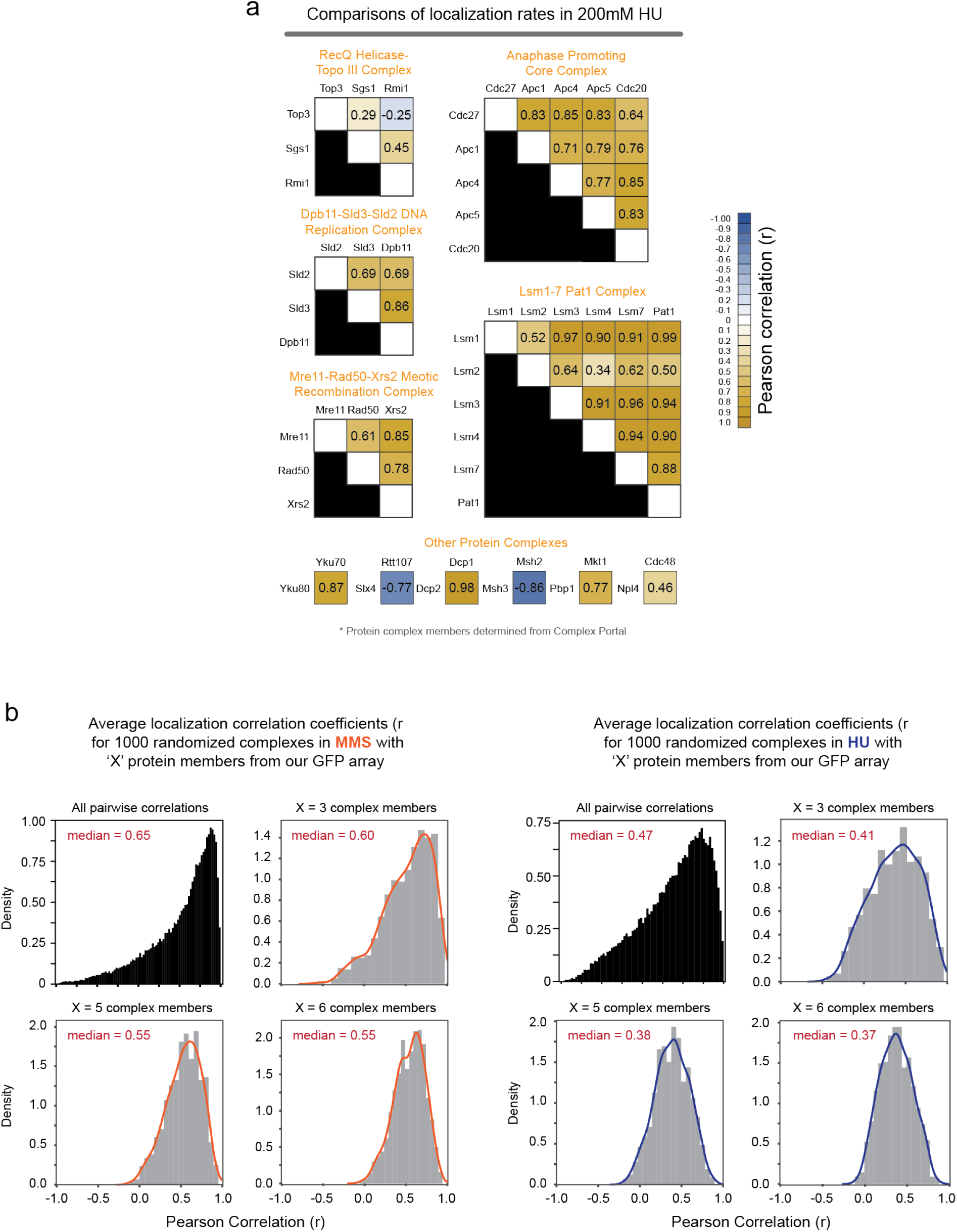
Correlation of protein localization kinetics among protein complex members. (a) Correlation matrices for 11 protein complexes represented in our set of proteins analyzed as identified by Complex Portal, displaying the pairwise Pearson correlation coefficients for each protein member of the complex, in 200mM HU conditions. (b) The distribution of all pairwise Pearson correlation coefficients between any two proteins, and the distribution of the average Pearson correlation coefficient for 1000 randomly generated protein complexes with the indicated number of proteins (X) taken from the proteins analyzed in this study.

## References

1. Almagro Armenteros, J.J., Sønderby, C.K., Sønderby, S.K., Nielsen, H., and Winther, O. (2017). DeepLoc: prediction of protein subcellular localization using deep learning. Bioinformatics 33, 3387–3395.

2. Breker, M., Gymrek, M., and Schuldiner, M. (2013). A novel single-cell screening platform reveals proteome plasticity during yeast stress responses. J. Cell Biol. 200, 839–850.

3. Breker, M., Gymrek, M., Moldavski, O., and Schuldiner, M. (2014). LoQAtE - Localization and Quantitation ATlas of the yeast proteomE. A new tool for multiparametric dissection of single-protein behavior in response to biological perturbations in yeast. Nucleic Acids Res. 42.

4. Causton, H.C., Ren, B., Koh, S.S., Harbison, C.T., Kanin, E., Jennings, E.G., Lee, T.I., True, H.L., Lander, E.S., and Young, R.A. (2001). Remodeling of yeast genome expression in response to environmental changes. Mol. Biol. Cell 12, 323–337.

5. Cejka, P., Plank, J.L., Bachrati, C.Z., Hickson, I.D., and Kowalczykowski, S.C. (2010). Rmi1 stimulates decatenation of double Holliday junctions during dissolution by Sgs1-Top3. Nat. Struct. Mol. Biol. 17, 1377–1382.

6. Chang, M., Bellaoui, M., Zhang, C., Desai, R., Morozov, P., Delgado-Cruzata, L., Rothstein, R., Freyer, G.A., Boone, C., and Brown, G.W. (2005). RMI1/NCE4, a suppressor of genome instability, encodes a member of the RecQ helicase/Topo III complex. EMBO J. 24, 2024–2033.

7. Chen, S.C., Zhao, T., Gordon, G.J., and Murphy, R.F. (2007b). Automated image analysis of protein localization in budding yeast. Bioinformatics 23.

8. Chong, Y.T., Koh, J.L.Y., Friesen, H., Duffy, K., Cox, M.J., Moses, A., Moffat, J., Boone, C., and Andrews, B.J. (2015). Yeast proteome dynamics from single cell imaging and automated analysis. Cell 161, 1413–1424.

9. Cremona, C.A., Sarangi, P., Yang, Y., Hang, L.E., Rahman, S., and Zhao, X. (2012). Extensive DNA Damage-Induced Sumoylation Contributes to Replication and Repair and Acts in Addition to the Mec1 Checkpoint. Mol. Cell 45, 422–432.

10. Dalal, C.K., Cai, L., Lin, Y., Rahbar, K., and Elowitz, M.B. (2014). Pulsatile dynamics in the yeast proteome. Curr. Biol. 24, 2189–2194.

11. Decker, C.J., and Parker, R. (2012). P-Bodies and Stress Granules: Possible Roles in the Control of Translation and mRNA Degradation. Cold Spring Harb. Perspect. Biol. 4.

12. Dénervaud, N., Becker, J., Delgado-Gonzalo, R., Damay, P., Rajkumar, A.S., Unser, M., Shore, D., Naef, F., and Maerkl, S.J. (2013). A chemostat array enables the spatio-temporal analysis of the yeast proteome. Proc. Natl. Acad. Sci. U. S. A. 110, 15842–15847.

13. Dohn, R., Xie, B., Back, R., Selewa, A., Eckart, H., Rao, P., and Basu, A. (2021). mDrop-seq: Massively parallel single-cell RNA-seq of Saccharomyces cerevisiae and Candida albicans 2 3. BioRxiv.

14. Gallina, I., Colding, C., Henriksen, P., Beli, P., Nakamura, K., Offman, J., Mathiasen, D.P., Silva, S., Hoffmann, E., Groth, A., et al. (2015). Cmr1/WDR76 defines a nuclear genotoxic stress body linking genome integrity and protein quality control. Nat. Commun. 6, 6533.

15. Gangloff, S., McDonald, J.P., Bendixen, C., Arthur, L., and Rothstein, R. (1994). The yeast type I topoisomerase Top3 interacts with Sgs1, a DNA helicase homolog: a potential eukaryotic reverse gyrase. Mol. Cell. Biol. 14, 8391–8398.

16. Gasch, A.P. (2007). The environmental stress response: a common yeast response to diverse environmental stresses. Yeast Stress Responses 1, 11–70.

17. Gasch, A.P., and Werner-Washburne, M. (2002). The genomics of yeast responses to environmental stress and starvation. Funct. Integr. Genomics 2, 181–192.

18. Gasch, A.P., Spellman, P.T., Kao, C.M., Carmel-Harel, O., Eisen, M.B., Storz, G., Botstein, D., and Brown, P.O. (2000). Genomic expression programs in the response of yeast cells to environmental changes. Mol. Biol. Cell 11, 4241–4257.

19. Gasch, A.P., Yu, F.B., Hose, J., Escalante, L.E., Place, M., Bacher, R., Kanbar, J., Ciobanu, D., Sandor, L., Grigoriev, I. V., et al. (2017). Single-cell RNA sequencing reveals intrinsic and extrinsic regulatory heterogeneity in yeast responding to stress. PLoS Biol. 15.

20. Grys, B.T., Lo, D.S., Sahin, N., Kraus, O.Z., Morris, Q., Boone, C., and Andrews, B.J. (2017). Machine learning and computer vision approaches for phenotypic profiling. J. Cell Biol. 216, 65–71.

21. Hammarström, P., Jiang, X., Hurshman, A.R., Powers, E.T., and Kelly, J.W. (2002). Sequence-dependent denaturation energetics: A major determinant in amyloid disease diversity. Proc. Natl. Acad. Sci. U. S. A. 99 Suppl 4, 16427–16432.

22. Huh, W.-K., Falvo, J. V., Gerke, L.C., Carroll, A.S., Howson, R.W., Weissman, J.S., O’Shea, E.K., and Shea, E.K.O. (2003). Global analysis of protein localization in budding yeast. Nature 425, 686–691.

23. Kehrein, K., Schilling, R., Möller-Hergt, B.V., Wurm, C.A., Jakobs, S., Lamkemeyer, T., Langer, T., and Ott, M. (2015a). Organization of Mitochondrial Gene Expression in Two Distinct Ribosome-Containing Assemblies. Cell Rep. 10, 843–853.

24. Kehrein, K., Möller-Hergt, B.V., and Ott, M. (2015b). The MIOREX complex - lean management of mitochondrial gene expression. Oncotarget 6, 16806–16807.

25. Koh, J.L.Y., Chong, Y.T., Friesen, H., Moses, A., Boone, C., Andrews, B.J., and Moffat, J. (2015). CY-CLoPs: A comprehensive database constructed from automated analysis of protein abundance and sub-cellular localization patterns in Saccharomyces cerevisiae. G3 Genes, Genomes, Genet. 5, 1223–1232.

26. Kraus, O.Z., Grys, B.T., Ba, J., Chong, Y., Frey, B.J., Boone, C., and Andrews, B.J. (2017). Automated analysis of high-content microscopy data with deep learning. Mol. Syst. Biol. 13, 924.

27. Kupiec, M., and Simchen, G. (1985). Arrest of the mitotic cell cycle and of meiosis in Saccharomyces cerevisiae by MMS. Mol. Gen. Genet. MGG 1985 2013 201, 558–564.

28. Levy, S.F., Ziv, N., and Siegal, M.L. (2012). Bet hedging in yeast by heterogeneous, age-correlated expression of a stress protectant. PLoS Biol. 10.

29. Lisby, M., Barlow, J.H., Burgess, R.C., and Rothstein, R. (2004). Choreography of the DNA damage response: spatiotemporal relationships among check-point and repair proteins. Cell 118, 699–713.

30. Liu, J., François, J.-M.M., and Capp, J.-P.P. (2019). Gene expression noise produces cell-to-cell heterogeneity in eukaryotic homologous recombination rate. Front. Genet. 10, 475.

31. Loll-Krippleber, R., and Brown, G.W. (2017). P-body proteins regulate transcriptional rewiring to promote DNA replication stress resistance. Nat. Commun. 8.

32. Lu, A.X., and Moses, A.M. (2016). An Unsupervised kNN Method to Systematically Detect Changes in Protein Localization in High-Throughput Microscopy Images. PLoS One 11, e0158712.

33. Lu, A.X., Chong, Y.T., Hsu, I.S., Strome, B., Handfield, L.F., Kraus, O., Andrews, B.J., and Moses, A.M. (2018). Integrating images from multiple microscopy screens reveals diverse patterns of change in the subcellular localization of proteins. Elife 7.

34. Mathew, V., Kumar, A., Jiang, Y.K., West, K., Tam, A.S., and Stirling, P.C. (2020). Cdc48 regulates intranuclear quality control sequestration of the Hsh155 splicing factor in budding yeast. J. Cell Sci. 133.

35. Mayer, C., Dimopoulos, S., Rudolf, F., and Stelling, J. (2013). Using cellX to quantify intracellular events. Curr. Protoc. Mol. Biol. 101, 1–20.

36. Mazumder, A., Pesudo, L.Q., McRee, S., Bathe, M., and Samson, L.D. (2013). Genome-wide single-cell-level screen for protein abundance and localization changes in response to DNA damage in S. cerevisiae. Nucleic Acids Res. 41, 9310–9324.

37. McQuin, C., Goodman, A., Chernyshev, V., Kamentsky, L., Cimini, B.A., Karhohs, K.W., Doan, M., Ding, L., Rafelski, S.M., Thirstrup, D., et al. (2018). CellProfiler 3.0: Next-generation image processing for biology. PLoS Biol. 16, 1–17.

38. Möller-Hergt, B.V., Carlström, A., Stephan, K., Imhof, A., and Ott, M. (2018). The ribosome receptors Mrx15 and Mba1 jointly organize cotranslational insertion and protein biogenesis in mitochondria. Mol. Biol. Cell 29, 2386–2396.

39. Newman, J.R.S., Ghaemmaghami, S., Ihmels, J., Breslow, D.K., Noble, M., DeRisi, J.L., and Weissman, J.S. (2006). Single-cell proteomic analysis of S. cerevisiae reveals the architecture of biological noise. Nature 441, 840–846.

40. Nissan, T., and Parker, R. (2008). Analyzing P-bodies in Saccharomyces cerevisiae. Methods Enzymol. 448, 507.

41. O’Connell, M.J., Walworth, N.C., and Carr, A.M. (2000). The G2-phase DNA-damage checkpoint. Trends Cell Biol. 10, 296–303.

42. Ouyang, W., Winsnes, C.F., Hjelmare, M., Cesnik, A.J., Åkesson, L., Xu, H., Sullivan, D.P., Dai, S., Lan, J., Jinmo, P., et al. (2019). Analysis of the Human Protein Atlas Image Classification competition. Nat. Methods 2019 1612 16, 1254–1261.

43. Pellicioli, A., Lee, S.E., Lucca, C., Foiani, M., and Haber, J.E. (2001). Regulation of Saccharomyces Rad53 checkpoint kinase during adaptation from DNA damage-induced G2/M arrest. Mol. Cell 7, 293–300.

44. Rendleman, J., Cheng, Z., Maity, S., Kastelic, N., Munschauer, M., Allgoewer, K., Teo, G., Zhang, Y.B.M., Lei, A., Parker, B., et al. (2018). New insights into the cellular temporal response to proteostatic stress. Elife 7.

45. Schmidt, G.W., Frey, O., and Rudolf, F. (2018). The CellClamper: A convenient microfluidic device for time-lapse imaging of yeast (Methods Mol Biol).

46. Sharif, H., and Conti, E. (2013). Architecture of the Lsm1-7-Pat1 Complex: A Conserved Assembly in Eukaryotic mRNA Turnover. Cell Rep. 5, 283–291.

47. Srikumar, T., Lewicki, M.C., Costanzo, M., Tkach, J.M., van Bakel, H., Tsui, K., Johnson, E.S., Brown, G.W., Andrews, B.J., Boone, C., et al. (2013). Global analysis of SUMO chain function reveals multiple roles in chromatin regulation. J. Cell Biol. 201, 145–163.

48. Teixeira, D., and Parker, R. (2007). Analysis of P-Body Assembly in Saccharomyces cerevisiae. Mol. Biol. Cell 18, 2274.

49. Tkach, J.M., Yimit, A., Lee, A.Y., Riffle, M., Costanzo, M., Jaschob, D., Hendry, J. a, Ou, J., Moffat, J., Boone, C., et al. (2012). Dissecting DNA damage response pathways by analysing protein localization and abundance changes during DNA replication stress. Nat. Cell Biol. 14, 966–976.

50. Torres, N.P., and Brown, G.W. (2015). A high-throughput confocal fluorescence microscopy platform to study DNA replication stress in yeast cells. Methods Mol. Biol. 1300.

51. Torres, N.P., Ho, B., and Brown, G.W. (2016). High-throughput fluorescence microscopic analysis of protein abundance and localization in budding yeast. Crit. Rev. Biochem. Mol. Biol. 51, 110–119.

52. Uphoff, S., Lord, N.D., Okumus, B., Potvin-Trottier, L., Sherratt, D.J., and Paulsson, J. (2016). Stochastic activation of a DNA damage response causes cell-to-cell mutation rate variation. 351, 1094–1097.

53. Vincent, M.S., and Uphoff, S. (2021). Cellular heterogeneity in DNA alkylation repair increases population genetic plasticity. Nucleic Acids Res. 49, 12320–12331.

54. Wilusz, C.J., and Wilusz, J. (2013). Lsm proteins and Hfq: Life at the 3 end. RNA Biol. 10, 592. Wu, L., Bachrati, C.Z., Ou, J., Xu, C., Yin, J., Chang, M., Wang, W., Li, L., Brown, G.W., and Hickson, I.D. (2006). BLAP75/RMI1 promotes the BLM-dependent dissolution of homologous recombination intermediates. Proc. Natl. Acad. Sci. U. S. A. 103, 4068–4073.

55. Xiao, M., Shen, X., and Pan, W. (2019). Application of deep convolutional neural networks in classification of protein subcellular localization with microscopy images. Genet. Epidemiol. 43, 330–341.

56. Yamamoto, A., Guacci, V., and Koshland, D. (1996). Pds1p, an inhibitor of anaphase in budding yeast, plays a critical role in the APC and checkpoint pathway(s). J. Cell Biol. 133, 99–110.

57. Yang, J., Bachrati, C.Z., Ou, J., Hickson, I.D., and Brown, G.W. (2010). Human topoisomerase IIIal-pha is a single-stranded DNA decatenase that is stimulated by BLM and RMI1. J. Biol. Chem. 285, 21426–21436.

